# Correspondence of Newcomb-Benford Number Law with Ecological Processes

**DOI:** 10.1101/2022.06.27.497806

**Authors:** Robert D. Davic

## Abstract

The Newcomb-Benford number law has been applied in the natural sciences for decades, with little ecological attention. Empirical data transformed into significant digits reveal statistical correspondence between the discrete Benford probability distribution and physical systems in dynamic equilibrium along a continuum of stability. Analytic methods are presented to detect this mathematical representation across multiple levels of ecological organization and spatial scale. Case studies demonstrate novel application to help identify bidirectional regime changes to alternative states of dynamic equilibrium. Widespread documentation of the surprising phenomenon is anticipated as ecologists revisit historic sets of random measurement data and design future sampling protocols. Controlled experiments with measurement variables that span multiple orders of magnitude would be well suited for future empirical and theoretical inquiry.

## Introduction

Organization in ecological systems results from negative and positive feedback interactions between organisms and their abiotic environment [1]. The complex mix of stochastic and non-random (natural selection) interactions generates interest to identify patterns that unify ecological processes at multiple hierarchical levels and spatial scales [2]. Pattern in empirical data has been attributed to ‘law-like’ mathematical representations [3-6], a phenomenon not well investigated in ecology [7].

The purpose of this note is to communicate statistical correspondence between a long-known probability distribution (the Newcomb-Benford law [NBL], Newcomb, 1881 [8], Benford, 1938 [9], Pinkham, 1961 [10]) with ecological processes related to complexity and stability. The Benford distribution is an empirical observation that the first significant digits (FSD) from certain sets of naturally occurring data (i.e., with minimal human intervention) follow a discrete logarithmic probability distribution (Figure 1, Table 1). The NBL has been applied in the natural sciences for decades [11-18], with little ecological attention. Potential ecological applications arise from reports the dynamic equilibrium of physical and social systems (defined *sensu* Thoms et al. [19] as long-term <u>balanced</u> fluctuations about short-term constantly changing system conditions) can be well described by the Benford FSD sequence (Table 2 [14-18]). The recursive theme is that interactions of system components that maintain structural stability [20] (i.e., persistence, constancy, resilience) can be represented by the informational distance between two discrete probability distributions, one from a FSD transformation of random measurements on system variables, the second from the regularity of FSD probabilities predicted by the Newcomb-Benford law (see Appendix Figures 1-4). As the distance between distributions increases in response to external perturbations, build-up of informational uncertainty can lead to a phase transition away from a balanced threshold of minimal entropy production [21-23]. Prigogine [24] identified ‘dissipative structures’ to denote self-organized dynamic systems with a propensity of minimal entropy production over time to maintain stability. The phenomenon is consistent with Ulanowicz’s ascendency view of ecological stability [25], defined as the ability of an ecosystem to “*remain within an arbitrarily-designated nominal range of behavior”* in response to changing environmental conditions.

**Table 1.**
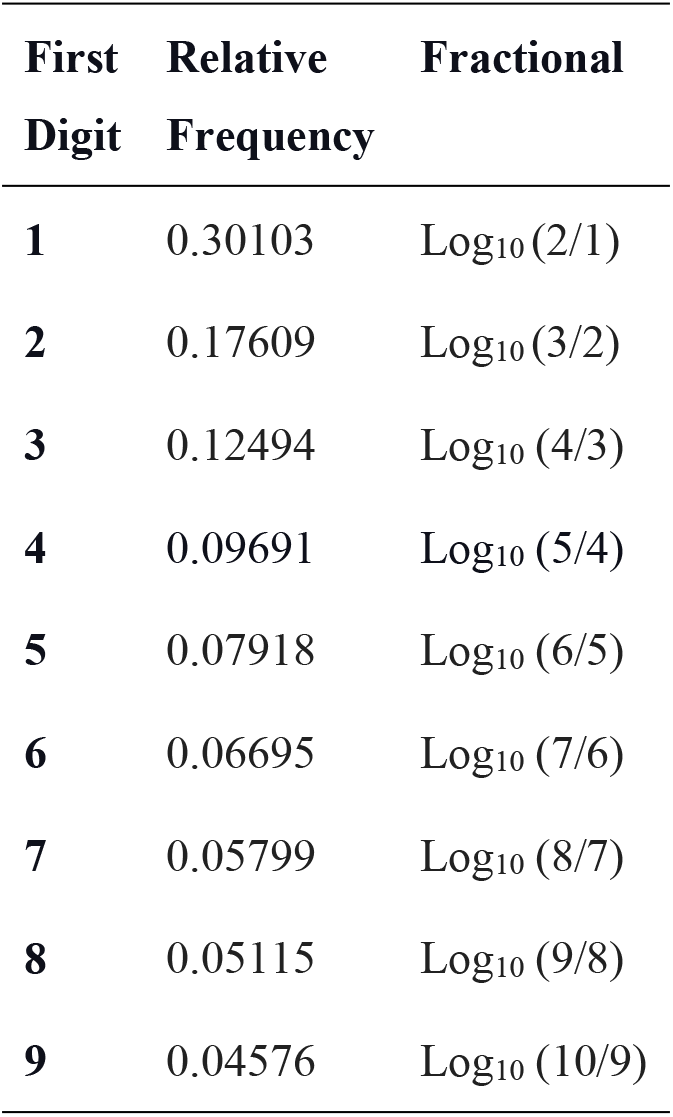
First significant digit relative frequencies for the Benford probability distribution.

| First<br>Digit | Relative<br>Frequency | Fractional |
| --- | --- | --- |
| 1 | 0.30103 | $\log_{10}(2/1)$ |
| 2 | 0.17609 | $\log_{10}(3/2)$ |
| 3 | 0.12494 | $\log_{10}(4/3)$ |
| 4 | 0.09691 | $\log_{10}(5/4)$ |
| 5 | 0.07918 | $\log_{10}(6/5)$ |
| 6 | 0.06695 | $\log_{10}(7/6)$ |
| 7 | 0.05799 | $\log_{10}(8/7)$ |
| 8 | 0.05115 | $\log_{10}(9/8)$ |
| 9 | 0.04576 | $\log_{10}(10/9)$ |

**Table 2.**
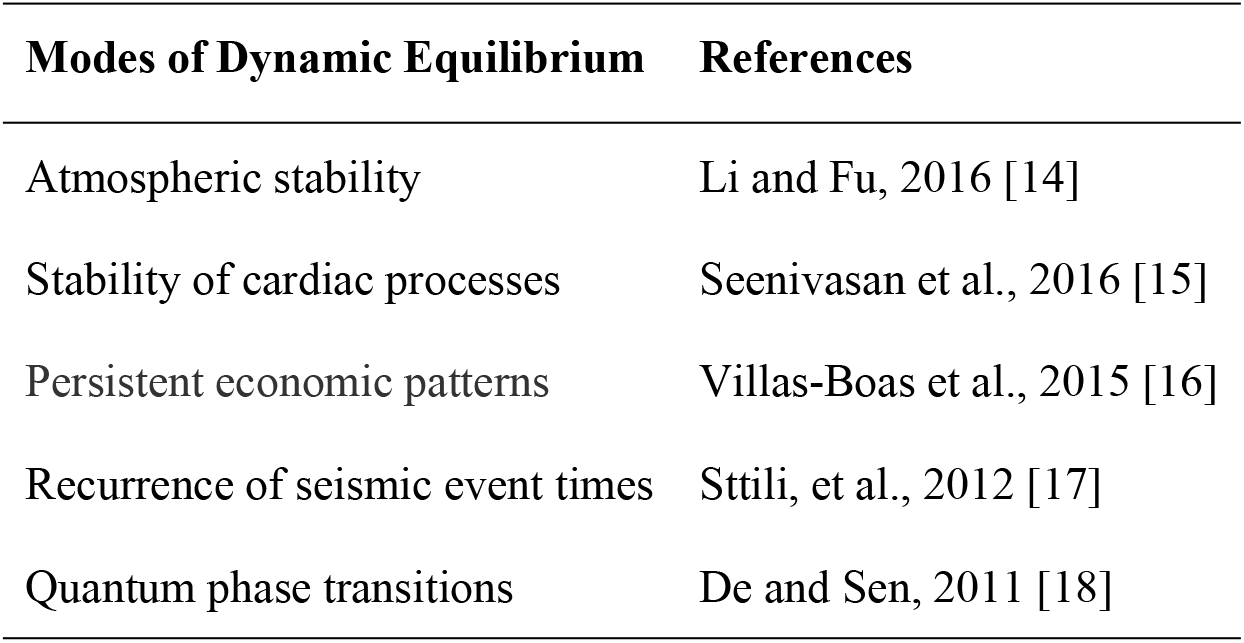
Physical and social systems in stable dynamic equilibrium that conform to the first significant digits frequencies of the Benford distribution.

**Figure 1.**
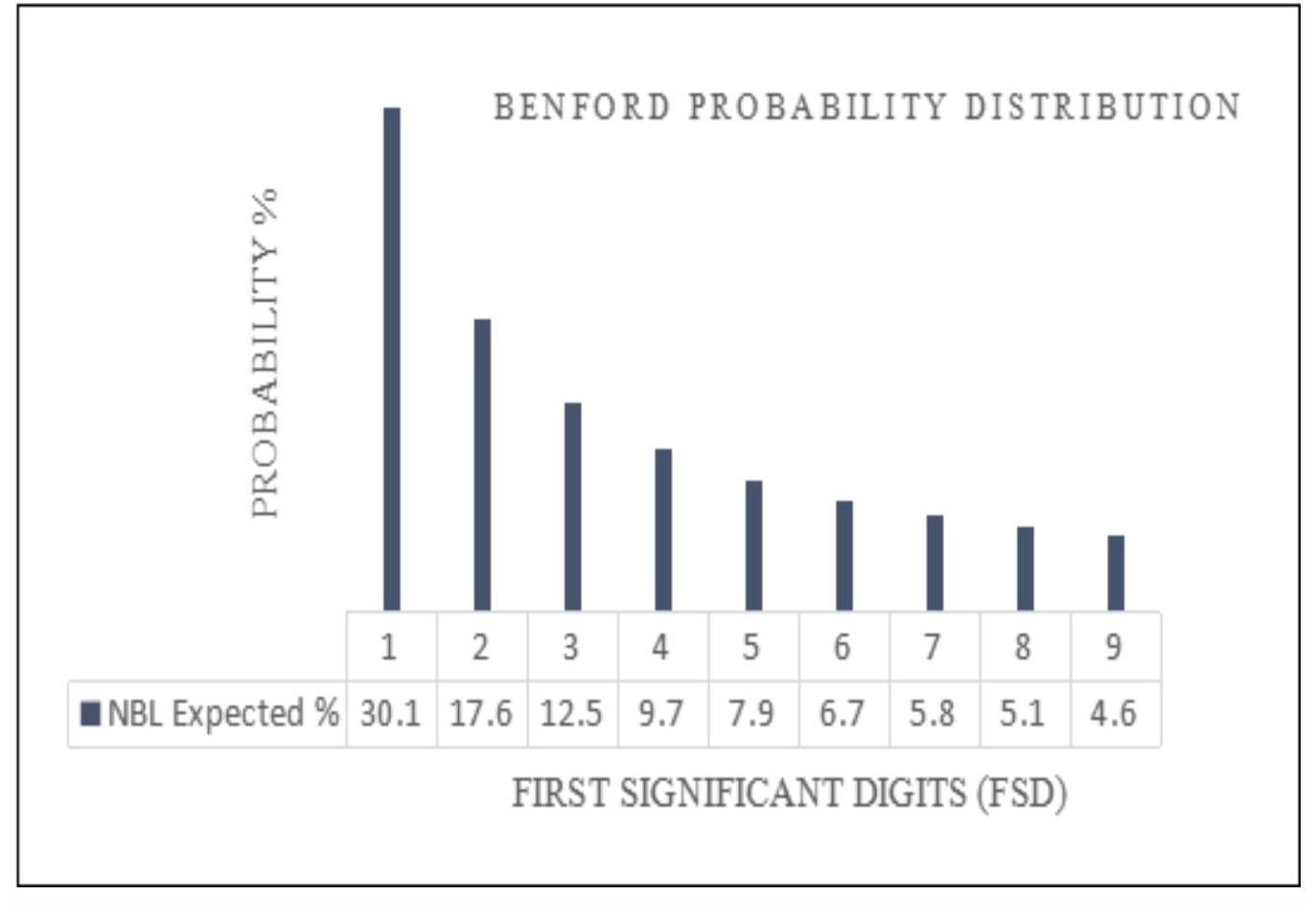
The discrete Benford probability distribution of first significant digits: P(*d*)=Log_10_ [1+(1/*d*)], where *d=1,2,…9* [8,9].

Given that dynamical ecological systems are open to short-term flux of matter, energy, and information, they also should adhere to reports of stability predicted by the first significant digit sequence of the NBL (Table 2). Anecdotal evidence comes from observations of seasonally predictable counts of lake algae that are well described by the Benford distribution [26,27], persistent gene sequences over evolutionary time scales that distinguish taxonomic categories above the species level [28], and annually consistent pollen counts from atmospheric samples [29]. Biotic communities at equilibrium have been associated with the log-normal distribution [30], which mathematically conforms with the Benford distribution [31]. Skewed Benford-like probability distributions (Figure 1) predict food-web stability [32] and have been suggested for use as an early warning signal of impending regime change for otherwise stable ecosystems [33].

After review of the Newcomb-Benford law, case reports of empirical data from the ecological literature are presented to test a proposition that naturally occurring ecological systems, reported to be progressing toward stable dynamic equilibrium, with minimal constraint violations related to collection of data, can be well represented by the Benford probability distribution.

### Modes of Dynamic Equilibrium References

Atmospheric stability Li and Fu, 2016 [14] Stability of cardiac processes Seenivasan et al., 2016 [15] Persistent economic patterns Villas-Boas et al., 2015 [16] Recurrence of seismic event times Sttili, et al., 2012 [17] Quantum phase transitions De and Sen, 2011 [18]

## Methods

### Background

The Newcomb-Benford law, also known as the first digit law and Benford’s law, is a mathematical oddity in the probability of first significant (non-zero) digits observed from certain sets of naturally occurring data (Figure 1). A Benford distribution emerges when a set of random measurements on system components i.e., (0.061, 0.14, 9.21, 20.2, 23.0, 723.0, 3345.0,…) are transformed into a compressed vector of FSD (6, 1, 9, 2, 2, 7, 3,…). The expected probability P(*d*) of leading digits (*d*) conforms to the logarithmic equation:xs

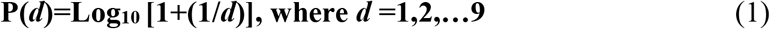

The leading digit #1 appears about 30% of the time, digit #9 near 5%, with a Log_10_ digit sequence (Table 1). First presented by Newcomb in 1881 [8], the phenomenon was rediscovered in 1938 by Benford [9] and empirically verified from 20,000+ measurements taken on sets of naturally occurring data compiled across multiple fields of study. The Benford distribution is uniquely both scale [10] and base [34] invariant and strongly correlated (R^2^=0.998, Table 1 data) with a power-law function over the first digit sequence (X1:9): Probability (Y)=0.3135 X ^(−0.8637)^. Increased conformity to law predictions is expected when data from multiple variables are aggregated. Additional information from analysis of second and third significant digits can improve the predictive power for some applications [35].

Hill [36] derived the Benford distribution to be a phenomenon of collecting random samples from multiple random distributions:*“if distributions are selected at random (in an unbiased way) and random samples are taken from each of the distributions, then the significant digit frequencies of the combined samples converge to Benford’s distribution though the individual distributions selected may not closely follow the law*.*”* The correspondence may not be intuitive to those that collect ecological sets of data. The NBL predicts that random measurements collected on organisms, for example the biomass of species nested within a herbivore functional group in a grassland habitat, will not yield a FSD sequence with as many biomass measurements that begin with 1, 2, 3 as 7, 8, 9 (e.g.,∼33.3% per triplet set), but instead the prior expectation is that the naturally occurring frequencies will be 60.2% and 15.5% respectively (Table 1).

Empirical support for the Benford phenomenon comes from multiple fields of study including weight of molecular compounds, numbers on economic spreadsheets, algae cell counts in lakes, earthquakes, and astronomic bright objects identified by the Fermi space telescope [12]. Mathematical functions of interest to ecologists that conform to the Benford distribution include: reciprocal density [37], exponential [38], log-normal [31], Laplace transform [39], Markov processes [40], and Poisson [41]. The most common application is from economics where deviation from a FSD sequence is used to detect the falsification of data on spreadsheets and tax returns [35,42]. Mathematical reviews include Berger and Hill, [43] and Miller [44]. A real-time online bibliography of Berger, Hill, and Rogers [45] provides 1000+ articles and books with theoretical and applied examples of the Newcomb-Benford law: http://www.benfordonline.net.

A variety of constraints (measurement requirements) associated with the collection and reporting of data are prerequisites for the emergence of the Benford distribution (Table 3). An increase in the number and magnitude of *constraint violations* increases the likelihood the set of data under investigation will not conform with NBL predictions. A review of literature identifies the following constraints associated with ecological applications: (1) the collection of data follows methods of random sampling such that each of the nine first digit categories has the same chance of occurrence for each measurement taken, with location of sample sites chosen to include habitat heterogeneity (i.e., such as riffles, runs, and deep pools in lotic systems); (2) the set of data span several orders of magnitude, ideally (10^6^) suggested by Fewster [46], but as few as (10^2^) reported here; (3) total sample size is sufficient to populate all FSD categories and appropriate for methods adopted for statistical inference; (4) the full range of data analyzed is not filtered within arbitrary limits; (5) the sampled habitat is naturally occurring with regional background levels of global environmental disturbance from climate change, and minimal direct anthropogenic perturbation (i.e., habitat alteration, chemical pollution); (6) tabulation of data is free from intentional human alteration; and (7) tabulated data are generated by non-additive mathematical operations such as multiplication, division, or numbers raised to integer powers [42,47], which are associated with multiple biotic processes.

**Table 3.**
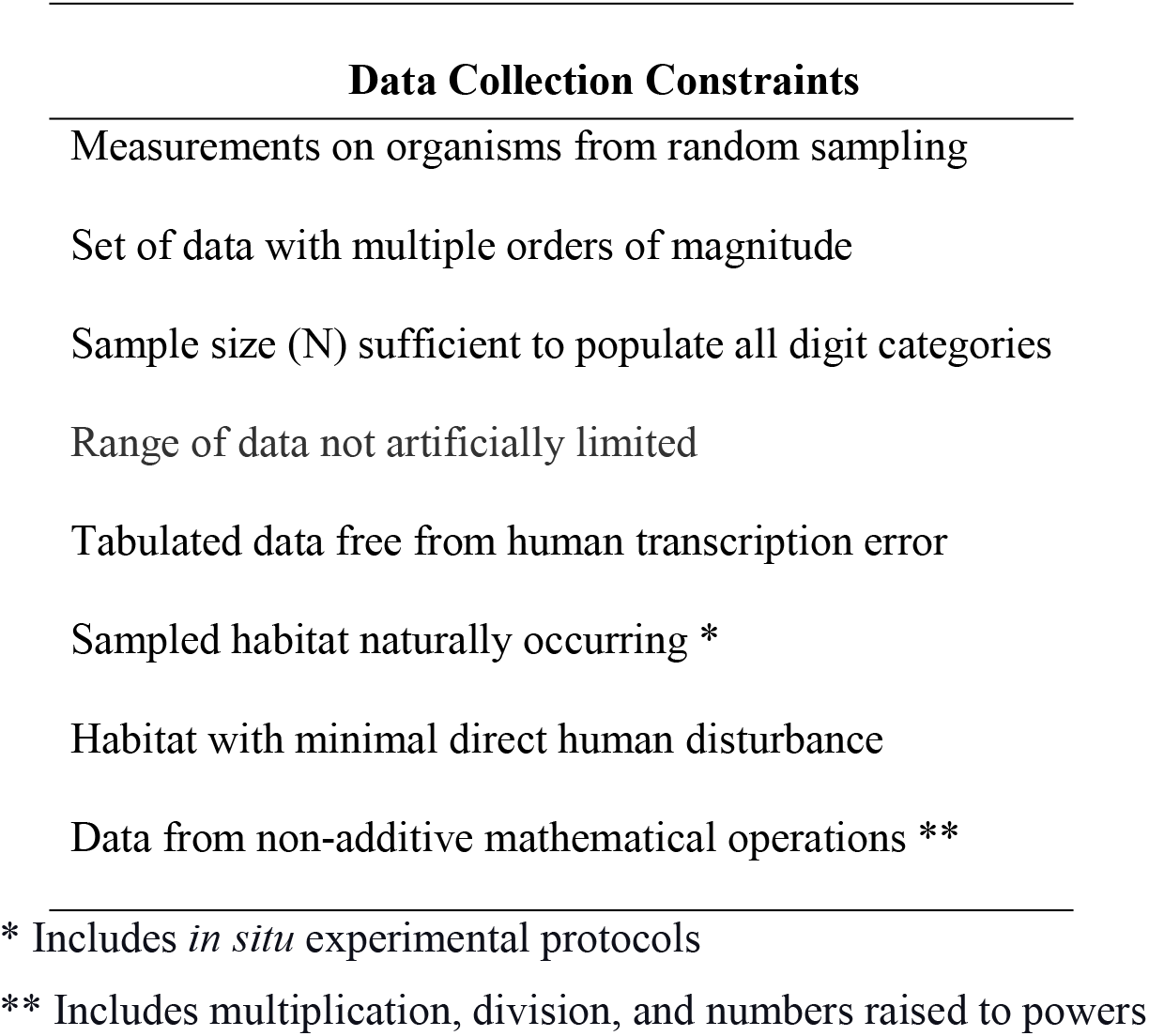
Data collection and reporting constraints associated with emergence of the Benford distribution.

| **Data Collection Constraints** |
| --- |
| Measurements on organisms from random sampling |
| Set of data with multiple orders of magnitude |
| Sample size (N) sufficient to populate all digit categories |
| Range of data not artificially limited |
| Tabulated data free from human transcription error |
| Sampled habitat naturally occurring * |
| Habitat with minimal direct human disturbance |
| Data from non-additive mathematical operations ** |

\* Includes *in situ* experimental protocols
\*\* Includes multiplication, division, and numbers raised to powers

The Benford FSD phenomenon has been demonstrated for a variety of ecological variables including counts of individuals and taxa, biomass, animal movements, and taxonomic classifications. Cell counts/colony of the cyanobacterium *Microcystis aeruginosa* collected from five lakes in Spain were well described by NBL predictions [25], as well as persistent communities of lake algae sampled over five years [26] and annually consistent pollen counts from atmospheric samples [28]. Campos et al. [48] observed that naturally assigned taxonomic categories of angiosperms conformed with NBL predictions, those artificially assigned did not. The number/ha of Red-Listed indicator species from 2707 conservation forest plots in Sweden visually conformed with the Benford distribution [49]. Pröger et al. [50] applied a Benford FSD analysis to telemetry data for wild red deer freely moving in natural habitats. Ninety one percent of 1132 weekly sets of data conformed with NBL predictions. Disturbance from human activity was suggested as a possible causal factor for non-conformance.

### Statistical Analysis

Empirical data from the ecological literature were analyzed to test a proposition that naturally occurring ecological systems, with minimal constraint violations related to collection of data, and reported to be progressing toward stable dynamic equilibrium, can be well represented by the Benford probability distribution. A weight-of-evidence analytic approach was adopted to quantify differences between microscopic and macroscopic attributes of observed *vs* expected FSD probability distributions.

#### Analytic Method #1

The following macroscopic null hypothesis (H_O_) was adopted: the measured nine FSD probability sequence is well represented by the sequence predicted by the Newcomb-Benford law, with alternative hypothesis (H_A_): the measured FSD sequence is not well represented. The (H_O_) was tested with a nine-dimensional Euclidean Distance formula, recommended for Benford applications by Coe and Gaines [51], as modified by Morrow [52] to allow for statistical inference:

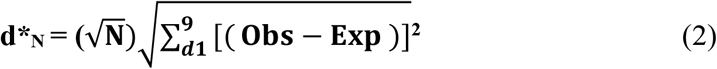

where *d* = digits (1:9), N = total sample size of observations, Obs = probability a random measurement *x* has a FSD = *d* for total sample size N, and Exp = Newcomb-Benford law expected FSD probabilities from Table 1 as Log_10_ (1+1/*d*).

A (H_O_) rejection range (alpha 0.10 d*_N_ = 1.22; alpha 0.05 d*_N_ = 1.33; alpha 0.01 d*_N_ = 1.57) is presented by Morrow derived from Monte Carlo simulation calibrated for sample size: N>80<500. Values of d*_N_ >1.33 reject the (H_O_). This narrow range of sample size is a limitation in the use of the Morrow (d*_N_) statistic, but should be sufficient for most ecological applications. Euclidean distance (>1.22<1.33) was adopted to serve as an early warning signal of impending regime change. Goodness of fit statistics for Benford applications can yield false negative errors at larger sample sizes [51,52,53]. Kossovsky [53] advocates use of sum of squared deviations (SSD) to quantify the magnitude of distance between measured data with NBL predictions. The SSD is calculated as percentages of [(Obs-Exp)]^2^ without consideration of the 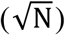 sample size operator (equation #2). Kossovsky proposes SSD>100 as a signal that measurement data are not well represented by the Benford probability distribution.

#### Analytic Method #2

Cohen [54] recommends calculation of *effect size* to estimate the magnitude of distance between two probability phenomena to complement interpretation of results from statistical inference. As relates to a Benford FSD analysis of data, a higher effect size is expected with an increase in the number and magnitude of constraint violations associated with collection of data (see Table 3). One measure of effect size that is independent of sample size and compliments information from Euclidean distance is the Cohen-W:

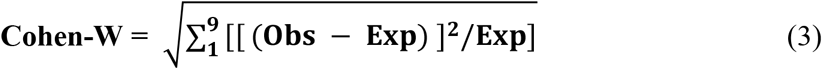

where Obs and Exp are the same as used to calculate Euclidean distance (d*_N_). Cohen [54] offers thresholds to identify weak, moderate, and strong effect sizes respectively (W=0.1, 0.3, 0.5). A Cohen-W>0.5 provides strong non-statistical inference of increased magnitude of constraint violations leading to non-conformity of NBL predictions. A Cohen-W range (>0.3<0.5) was adopted to serve as an early warning signal of impending regime change resulting from effects on skewness related with constraint violations.

#### Analytic Method #3

Microscopic multivariate volatility of FSD frequencies (e.g., entropy production) was estimated from the Pearson Residual (PR) formula presented by Sharpe [55]:

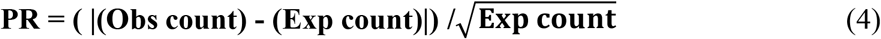

where Obs = measured FSD count frequency, and Exp = predicted FSD frequency based on total sample size N (e.g., from Table 1, the Exp counts for FSD #1= 0.30103*N). A PR value >1.96 (Z-Test, p = 0.05) for any FSD category was adopted as an early warning signal of impending regime change related to interaction effects of organisms with each other and their abiotic environment.

## Results

Analytic results for four case reports are presented in Appendix A and summarized in Table 4. The weight of evidence supports a proposition that naturally occurring ecological systems progressing toward a state of stable dynamic equilibrium can be well represented by the first significant digit sequence of the Newcomb-Benford law. The phenomenon was demonstrated at multiple levels of organization and spatial scales: (1) annual functional group biomass of below ground food-webs during developmental stages of succession, (2) counts of diatom cells on glass slides progressing toward dynamic equilibrium of taxa richness during island colonization, (3) constancy in abundances of meta-populations of woodland salamanders over a large geographic area, and (4) biomass of isolated meta-communities of lotic freshwater fish with persistent biotic integrity. Inclusive conformity with FSD predictions of the the Benford distribution, for five case report comparisons, was within analytical ranges far away from a threshold limit of minimal entropy production (Table 4): Euclidean distance (d*_N_= 0.638-0.783), Cohen-W effect size (0.171-0.250), Max-PR entropy (1.13-1.87), and Kossovsky SSD (21.2-63.9). Results from two case studies highlight the importance of including analytic test results from Pearson Residuals and Cohen-W effect size to help detect signals of impending regime change to compliment information from goodness-of-fit statistical inference.

**Table 4.**
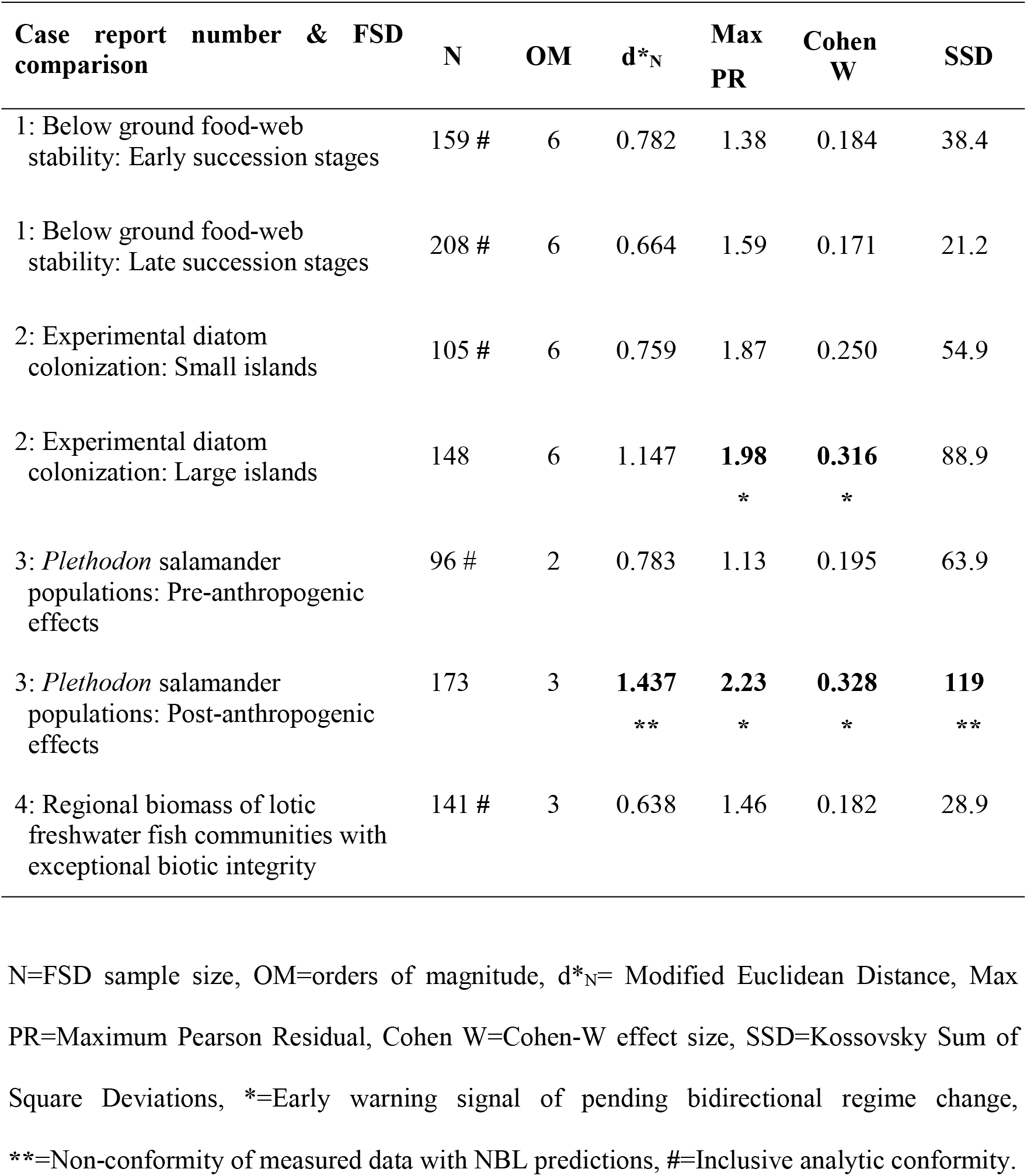
Summary of analytic test results from Appendix A case reports.

| Case report number & FSD comparison | N | OM | d*N | Max PR | Cohen W | SSD |
| --- | --- | --- | --- | --- | --- | --- |
| 1: Below ground food-web stability: Early succession stages | 159 # | 6 | 0.782 | 1.38 | 0.184 | 38.4 |
| 1: Below ground food-web stability: Late succession stages | 208 # | 6 | 0.664 | 1.59 | 0.171 | 21.2 |
| 2: Experimental diatom colonization: Small islands | 105 # | 6 | 0.759 | 1.87 | 0.250 | 54.9 |
| 2: Experimental diatom colonization: Large islands | 148 | 6 | 1.147 | **1.98**<br>* | **0.316**<br>* | 88.9 |
| 3: *Plethodon* salamander populations: Pre-anthropogenic effects | 96 # | 2 | 0.783 | 1.13 | 0.195 | 63.9 |
| 3: *Plethodon* salamander populations: Post-anthropogenic effects | 173 | 3 | **1.437**<br>** | **2.23**<br>* | **0.328**<br>* | **119**<br>** |
| 4: Regional biomass of lotic freshwater fish communities with exceptional biotic integrity | 141 # | 3 | 0.638 | 1.46 | 0.182 | 28.9 |

N=FSD sample size, OM=orders of magnitude, d\*N= Modified Euclidean Distance, Max PR=Maximum Pearson Residual, Cohen W=Cohen-W effect size, SSD=Kossovsky Sum of Square Deviations, \*=Early warning signal of pending bidirectional regime change, \*\*=Non-conformity of measured data with NBL predictions, #=Inclusive analytic conformity.

## Discussion

MacArthur [57] suggested the science of ecology is in large part a search for repeating mathematical patterns in nature. The Fibonacci pattern (1,1,2,3,5,8,13,21…), first applied to explain breeding in rabbits, is associated with the phenotypic expression of amino acids during embryonic development [58]. Margalef [59] equated the cybernetic stability of ecosystems to the ‘constancy of numbers’. Data presented here support a hypothesis that ecological systems progressing toward a threshold of stable dynamic equilibrium can be well represented by constancy in the sequence of first significant digits predicted by the Newcomb-Benford number law (Figure 1). The results concur with similar observations for multiple physical and social systems (Table 2), mathematical constants of nature [56], and anecdotal biotic observations [26-29,48-50].

Skewed probability distributions have been suggested to detect ecological systems that depart from stability thresholds [33]. Case reports suggest a hypothesis that changes in skewness characteristics of the Benford FSD distribution can be used as a bidirectional early warning signal of regime change. Results from a colonization experiment (case report #2) document the natural progression of a diatom community towards increasing dynamic equilibrium of species richness as predicted by theory of island biogeography, whereas analysis of long-term observations of persistent meta-populations of woodland salamanders (case report #3) reveal regime change away from stable species abundances in response to human-influenced environmental perturbation.

An important caveat for ecological application is that not all biotic systems in stable dynamic equilibrium that are well described by NBL predictions will necessarily be of beneficial use to humans. Examples include lentic habitats with seasonally stable blooms of cyanobacteria that can reach limits of toxicity to wildlife, and biotic communities with dominant and stable populations of exotic species that alter community biotic integrity and well-being. Research directed toward a mix of sample variables is suggested to partition the detection of the Benford distribution with and without the presence of non-beneficial populations.

A Benford FSD analysis of random measurement data presents a variety of theoretical and empirical lines of inquiry for future ecological research: (1) demonstration of sample size effects on NBL predictions in the range N>80<500 at multiple levels of organization and spatial scale, (2) expansion of the list of measurement variables, (3) identification of alternative statistical methods for early identification of regime shifts, (4) experiments investigating data collection constraints associated with human perturbation of environmental conditions, (5) analyses of second and third significant digits to improve predictive power, (6) comparisons of random sampling methods, (7) sets of big-data created via remote sensing [60], and (8) a FSD analysis using metabolism of detritus as a measurement variable, suggested by Wetzel [61] to be an integral driver of stability for both aquatic and terrestrial ecosystems. Further investigation also is warranted concerning the mechanisms by which the Newcomb-Benford mathematical representation is compatible with ecological concepts related to scaling laws, including Taylor’s power law [62], MacArthur’s broken stick model of niche appointments [63], maximum information entropy [64,65], effective complexity [66], ecosystem sustainability [67], and cybernetic Markovian processes [68,40].

Consensual agreement on the measurement of ecological stability remains elusive. The prevailing view of Landi et al. [69] is that no single system parameter, such as species diversity, can aggregate all possible interpretations of stability. Ives and Carpenter [70] argue for a holistic approach: “*it will be more profitable to study stability comprehensively, including diversity as only one of the possible factors that affect ecosystem responses to environmental change*.*”* A Benford FSD analysis of measurement data can serve to quantify the comprehensive stability of complex adaptive systems. Following Ulanowicz’s ascendency view of stable ecological development [25], as measurements collected on organisms transition from one microscopic FSD category to another (i.e., a measurement at time *x* of 0.19 transforms to 0.20 at *x*+1 and populates a higher first digit category), the macroscopic sequence of FSD probabilities remains within a threshold range of minimal entropy production. Comprehensive system stability (*sensu* Ives and Carpenter) emerges as information gained from the FSD probabilities of measurement *variables* (i.e., system properties that vary over time from one individual to another), with additional data from statistical *parameters*, such as species diversity, applied to dynamical modelling to monitor factors that affect responses of structural stability to environmental change.

Wilson [71] proposed that future research in ecology be directed towards a better understanding of “*transformations that reassemble components in ways that retain system properties holistically,”* with future success measured by the ability of such transformations to identify emergent phenomena that are useful. A simple Benford FSD transformation of arithmetic data randomly collected on organisms, reassembled into a mathematical representation of first significant digit categories of logarithmic information, affords one holistic approach forward to investigate this consilience goal.

## Conclusion

Reviewed empirical data support a hypothesis that complex adapted systems trending toward a threshold of minimal entropy production and stable dynamic equilibrium can be well represented by the Benford probability distribution of first significant digits. Analytic methods using multidimensional Euclidean distance and Cohen-W effect size magnitude are presented to detect the surprising correspondence at multiple levels of ecological hierarchical organization and spatial scales. Case studies demonstrate that the Benford distribution can serve as a bidirectional early warning signal of regime change in response to both naturally occurring and anthropogenic modified environmental conditions. Widespread documentation of the mathematical representation is expected as ecologists revisit historic data and design sampling protocols with attention to constraints associated with collection of data. Experiments using conserved variables that span multiple orders of magnitude, for populations with a short generation time, would be well suited for future empirical and theoretical inquiry. A simple transformation of an arithmetic set of random measurement data into a vector of first significant digits, nested into discrete categories of logarithmic probabilities, offers novel potential to monitor the effective comprehensive stability and complexity of complex adaptive systems over ecological and evolutionary lengths of time.

## Data Availability

The availability of data cited in Appendix A case reports is included in the text.

## Acknowledgments

The author appreciates discussions with Don Lemons and Alex Kossovsky about methods of data analysis during preparation of the manuscript and comments on the manuscript. John Morrow granted permission for use of his modified Euclidean distance formula. A preprint version of this manuscript is displayed at the Cold Spring Harbor Laboratory ‘bioRxiv’ server [83].

## APPENDIX A: CASE REPORTS

### Case Report #1: Food-web stability during succession

#### Background

Neutel et al., [72] sampled two below ground food-webs representing four developmental stages of succession at the Schiermonnikoog [SCH] island national park and Hulschorsterzand [HUL] mainland nature reserve, Netherlands. Food-web stability was quantified from dynamic models related to three omnivorous feedback loops. Non-random predator-prey interactions within these loops were reported to play an important role preserving food-web stability at both sample locations as productivity and functional group complexity increased during succession.

Mean functional group biomass data (gms C ha^-1^ cm depth^-1^) are provided in online supplemental Table 1 of Neutel et al. [doi:10.1038/nature06154]. Soil samples were randomly collected in upper soil layers with four replications per development stage, three times, to establish yearly mean biomass statistics. Measurements span six orders of magnitude at both sample locations with sufficient sample size to test conformity with NBL predictions when data are combined for both SCH and HUL locations, and segregated into early (1,2) and late (3,4) developmental stages. Minimal anthropogenic disturbance is expected for the two natural areas.

#### Results and discussion

Functional group biomass counts at combined SCH and HUL locations provide insufficient weight of evidence to reject the tested (H_O_) of similarity between observed *vs* NBL expected FSD probability distributions (Appendix Table 1). Pearson Residuals (max<1.96) and Cohen-W effect sizes (max <0.18) agree with results from Euclidean distance (max d*n<0.782, p>0.10) that mean annual functional group biomass is well described by the Benford distribution, within a nominal range of uncertainty, at both early and late developmental stages of succession (Appendix Table 1). The Benford FSD analysis agrees with findings of Neutel et al. that the SCH and HUL food-webs have maintained stable dynamic equilibrium over time (Appendix Figure 1). No evidence of pending regime change was observed for combined SCH and HUL locations. As documented by Neutel et al., [72] an increase in FSD functional group diversity over time (N 159 to 208, Appendix Table 1) did not lead to a decrease in comprehensive food-web stability as would be predicted by purely random models of species interactions.

Altenia et al. [73] revisited data of Neutel et al. to investigate effects of weighted connectence of biomass flux rates on food-web stability. The study was motivated in part by reports that species interactions in ecosystems are not evenly distributed in space and time, but instead follow skewed probability patterns, such as observed in Appendix Figure 1. A Benford FSD analysis of biomass flux rate data provides a novel approach to investigate Altenia et al.’s finding that food-web stability has increased over time at the individual SCH and HUL sample locations, a trend suggested at combined sample locations (Appendix Table 1).

**Appendix Table 1.** Observed first significant digit counts of annual biomass for fifteen functional groups *vs* counts expected by the Benford probability distribution. Counts from combined SCH and HUL sample locations during early and late stages of food-web development. Raw data from Neutel et al., Supp. Table 1 [72].

**EARLY SUCCESSION (DEVELOPMENT STAGES 1 & 2)**
| **First Digits** | **1** | **2** | **3** | **4** | **5** | **6** | **7** | **8** | **9** |
| --- | --- | --- | --- | --- | --- | --- | --- | --- | --- |
| Observed counts | 42 | 29 | 26 | 16 | 12 | 9 | 13 | 7 | 5 |
| NBL Expected Counts | 47.9 | 28.0 | 19.9 | 15.4 | 12.6 | 10.6 | 9.2 | 8.1 | 7.3 |
| Pearson Residuals | 0.85 | 0.19 | 1.38 | 0.15 | 0.17 | 0.50 | 1.24 | 0.40 | 0.84 |
| Observed Probability % | 26.4 | 18.2 | 16.4 | 10.1 | 7.5 | 5.7 | 8.2 | 4.4 | 3.1 |
| Expected Probability % | 30.1 | 17.6 | 12.5 | 9.7 | 7.9 | 6.7 | 5.8 | 5.1 | 4.6 |

**LATE SUCCESSION (DEVELOPMENT STAGES 3&4)**
| **First Digits** | **1** | **2** | **3** | **4** | **5** | **6** | **7** | **8** | **9** |
| --- | --- | --- | --- | --- | --- | --- | --- | --- | --- |
| Observed counts | 64 | 36 | 28 | 22 | 10 | 19 | 13 | 7 | 9 |
| NBL Expected Counts | 62.6 | 36.6 | 26.0 | 20.2 | 16.5 | 13.9 | 12.1 | 10.6 | 9.5 |
| Pearson Residuals | 0.18 | 0.10 | 0.39 | 0.41 | 1.59 | 1.36 | 0.27 | 1.12 | 0.17 |
| Observed Probability % | 30.8 | 17.3 | 13.5 | 10.6 | 4.8 | 9.1 | 6.2 | 3.4 | 4.3 |
| Expected Probability % | 30.1 | 17.6 | 12.5 | 9.7 | 7.9 | 6.7 | 5.8 | 5.1 | 4.6 |

FSD sample size N = 208. Max Pearson Residual<1.96; Euclidean distance d\*n = 0.664 (p>0.10); Cohen-W = 0.171 (95% CI: 0.127-0.216); Kossovsky SSD = 21.2

**Appendix Figure 1.**
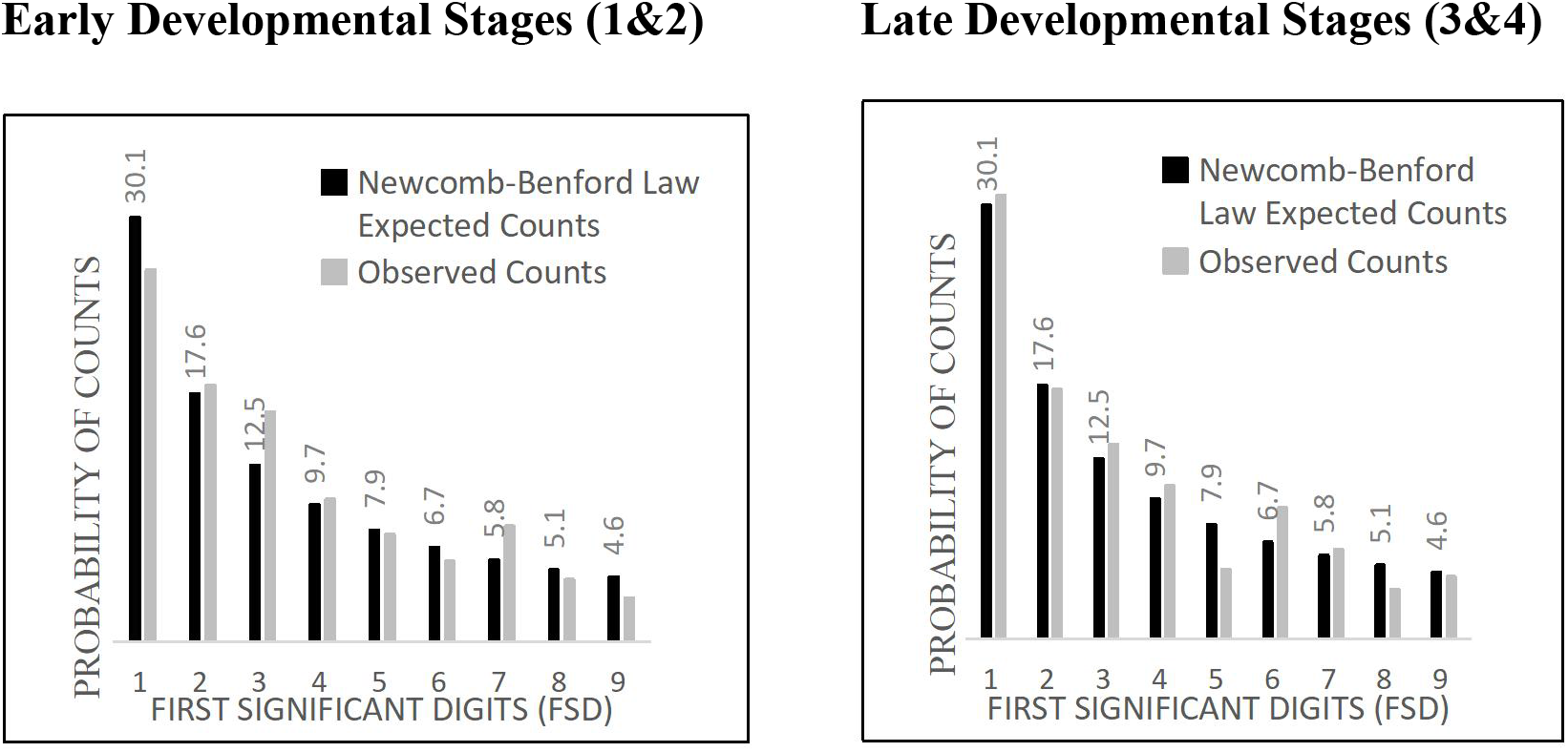
Comparison between FSD counts expected by Newcomb-Benford law and the probability distribution of observed counts for annual biomass of fifteen functional groups at combined SCH and HUL sample locations [72]. Graphs of data from Appendix Table 1.

### Case Report #2: Dynamic equilibrium of diatom community colonization

#### Background

MacArthur and Wilson [74], their Table 6, provide diatom cell counts for a colonization experiment conducted by Ruth Patrick at Roxborough Spring, Pennsylvania. Over a span of two weeks, diatom species colonized glass slides representing two size ‘islands’ (12 & 25 mm^2^). Replicate samples are reported as experiments 1&3 and 2&4. At experiment end, a diverse community of 54 diatom taxa and >600,000 diatom cell counts were recorded. MacArthur and Wilson cited the Patrick data in support of their equilibrium model of island biogeography, which predicts the ecological process of diatom colonization was progressing toward a threshold of stable dynamic equilibrium of species richness in response to stabilizing selection effects in rates of species immigration and extinction.

The diatom count data of Patrick meet sampling constraints required to test correspondence with NBL predictions (Table 3). The set of data spans six orders of magnitude with sufficient sample size to test conformity with the Benford distribution when data from replicated experiments for same sized islands are combined over the two week span of colonization. The experimental spring was described elsewhere by Patrick [75] as perennial flowing and free from human disturbance.

#### Results and discussion

Diatom cell counts for *smaller islands* provide insufficient analytic evidence to reject the tested (H_O_) of similarity between observed *vs* expected FSD probability distributions (Appendix Table 2 & Figure 2). No early warning evidence of pending regime change is evident (max PR<1.96; Cohen-W<0.3; d*n<1.22). Test results support an interpretation the diatom community on the smaller islands attained a state of stable dynamic equilibrium. Supporting evidence for this hypothesis comes from independent research at Roxborough Spring by Patrick [75] where constancy of diatom taxa richness was recorded after a two week span of colonization for an eight week experiment.

In contrast to results for smaller sized islands, analytic data for *larger islands* reveal higher levels of FSD volatility (Appendix Table 2), with both Pearson Residual (PR>1.96) and Cohen-W effect size (W = 0.316, 95% CI:0.250-0.381) within an early warning range of regime change. Theory of island biogeography predicts the increased volatility in response to effects of species interactions related to competition and taxa extinctions, a hypothesis open to experimental investigation by extending the colonization time for larger size islands to eight weeks. Results highlight the importance of including measures of Pearson Residual volatility and magnitude of Cohen effect size to complement statistical inference from calculation of Euclidean distance to detect marginally stable systems trending toward future bidirectional regime change (Appendix Table 2). Future research is suggested using protocols of Patrick [75] to increase sample size and island replicates to monitor time-series trends of FSD frequencies, as well as controlled experiments that manipulate data collection constraints related to human bias and alteration of habitat conditions.

**Appendix Table 2.** Observed first digit cell counts *vs* Newcomb-Benford law expected counts for diatom colonization experiments (MacArthur and Wilson [74], their Table 6).

**EXPERIMENTS 1&3 (12 mm<sup>2</sup> slides, small ‘islands’)**
| **Combined One & Two Week Exposure** |  |  |  |  |  |  |  |  |  |
| --- | --- | --- | --- | --- | --- | --- | --- | --- | --- |
| **First Digits** | **1** | **2** | **3** | **4** | **5** | **6** | **7** | **8** | **9** |
| Observed counts | 32 | 15 | 9 | 11 | 7 | 12 | 6 | 7 | 6 |
| NBL Expected Counts | 31.6 | 18.5 | 13.1 | 10.2 | 8.3 | 7.0 | 6.1 | 5.4 | 4.8 |
| Pearson Residuals | 0.07 | 0.81 | 1.14 | 0.26 | 0.46 | 1.87 | 0.04 | 0.70 | 0.55 |
| Observed Probability % | 30.5 | 14.3 | 8.6 | 10.5 | 6.7 | 11.4 | 5.7 | 6.7 | 5.7 |
| Expected Probability % | 30.1 | 17.6 | 12.5 | 9.7 | 7.9 | 6.7 | 5.8 | 5.1 | 4.6 |

**EXPERIMENTS 2&4 (25 mm<sup>2</sup> slides, large ‘islands’)**
| **Combined One & Two Week Exposure** |  |  |  |  |  |  |  |  |  |
| --- | --- | --- | --- | --- | --- | --- | --- | --- | --- |
| **First Digits** | **1** | **2** | **3** | **4** | **5** | **6** | **7** | **8** | **9** |
| Observed counts | 48 | 26 | 27 | 19 | 5 | 7 | 4 | 4 | 8 |
| NBL Expected Counts | 44.6 | 26.1 | 18.5 | 14.3 | 11.7 | 9.9 | 8.6 | 7.6 | 6.8 |
| Pearson Residuals | 0.52 | 0.01 | **1.98** | 1.23 | 1.96 | 0.92 | 1.56 | 1.30 | 0.47 |
| Observed Probability % | 32.4 | 17.6 | 18.2 | 12.8 | 3.4 | 4.7 | 2.7 | 2.7 | 5.4 |
| Expected Probability % | 30.1 | 17.6 | 12.5 | 9.7 | 7.9 | 6.7 | 5.8 | 5.1 | 4.6 |

FSD sample size N = 148. Max Pearson Residual>1.96; Euclidean distance d\*n = 1.147 (p>0.10); Cohen-W = 0.316 (95% CI: 0.250-0.381); Kossovsky SSD = 88.9.

**Appendix Figure 2.**
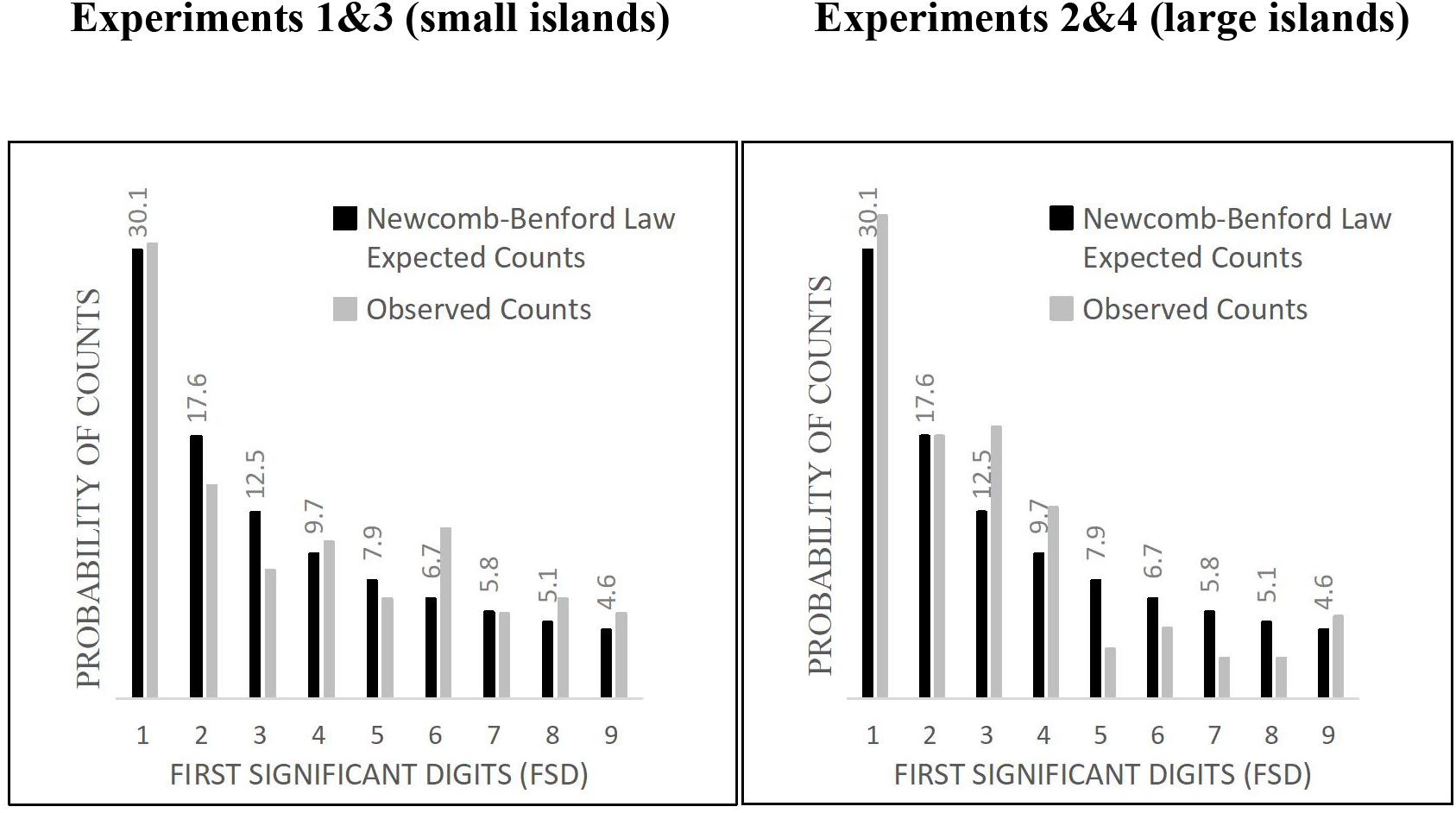
Comparison between FSD counts expected by Newcomb-Benford law and the probability distribution of observed counts for diatom cells colonizing islands at small and large spatial scales [74]. Graphs of data from Appendix Table 2.

### Case Study #3. Regional decline of stable amphibian populations

#### Background

Published databases suitable for a Benford FSD analysis with random samples collected before and after human disturbance of natural habitats, that also meet required data collection constraints (Table 3), are not common in the ecological literature. One example known to the author is that of Highton [76] for visual encounters of woodland salamanders (genus *Plethodon*) that span five decades. From 1957-1999 Highton and colleagues recorded observation of individual salamanders from 38 species of the genus *Plethodon* from 127 sample locations in forested habitats of Eastern United States. Pre-1990 surveys revealed long-term constancy of salamander abundance for 205 *Plethodon* populations (1957-1988). Post-1990 surveys (1991-1999) revealed decline in many of previously stable populations, with a statistically significant reduction in mean observations/site visit from 8.77 to 3.65 (Sign Test, χ^2^ = 117, p < 0.001). At 32 previously stable locations, *Plethodon* salamander species were extirpated. Although the causes for salamander declines were not investigated, logging was observed as a possible factor at multiple locations. Fungal infections, warming of forest temperatures from climate change, and changes in soil pH from acid rain are other suspected long-term stresses for woodland adapted salamanders.

The mean number of salamanders observed/site for Pre-1990 and Post 1990 are given in Table 8-1 of Highton [76]. The data span two and three orders of magnitude respectively. With the exception of the observed human habitat alteration for the Post-1990 surveys, the data meet sampling constraints required to test correspondence with NBL predictions (Table 3). Data were normalized to 1-6 site visits per observer, the maximum number reported for post-1990 surveys. This normalization process resulted in sample sizes N=96 (pre-1990) and N=173 (post-1990) to allow for statistical analysis.

#### Results and Discussion

Pre-1990 observations of salamander encounters provide insufficient weight of evidence to reject the tested (H_O_) of similarity between observed *vs* expected FSD probability distributions (Appendix Table 3 & Figure 3). Pearson Residuals (max<1.96) and Cohen-W effect size magnitudes (max<0.2) agree with results from statistical inference (d*n=0.783, p>0.10) that the regional abundances of 38 *Plethodon* salamander species were at stable dynamic equilibrium prior to 1988 surveys. This interpretation from the Benford FSD analysis matches pre-1988 observations of Highton [76] of “*consistent patterns of salamander surface activity from year to year”* at the 127 sample locations.

In contrast with Pre-1990 observations, Post-1990 test results reject the tested (H_O_). Signals of the regime change are presented for all analytical test methods (Pearson Residuals>1.96, Euclidean distance d*n = 1.437 (p<0.05), Cohen-W = 0.328 (95%CI 0.261-0.394). The Kossovsky SSD (119.4) adds support to an interpretation that the Post-1990 data are not well described by the Benford distribution (Appendix Table 3 & Figure 3). Results from the Benford FSD analysis concur with observation of Highton that widespread populations of *Plethodon* salamander species in the eastern United States from 1990 to 1999 were in various local modalities of unstable dynamic equilibrium compared to pre-1988 observations.

Post-1999 surveys on *Plethodon* species within the geographic range sampled by Highton by Caruso and Lips [77] and Caruso et al. [78] confirm the continuation of salamander declines, in particular at sample locations dominated by larger-bodied *Plethodon* species. Future investigation of measurement variables associated with gene flow among geographically isolated populations and allocation of energy to growth is suggested to investigate Benford FSD predictions of stability for those remaining large-bodied *Plethodon* populations under greatest selective pressure for extirpation.

**Appendix Table 3.** Observed first digit counts of mean number of *Plethodon* salamanders per site visit *vs* Newcomb-Benford law expected counts for stable (pre-1990) and declining (post-1990) populations from Eastern United States. (Highton [76], his Table 8-1). Analyzed data normalized to 1-6 site visits.

**Stable Pre-1990 Salamander Counts**
| **First Digits** | **1** | **2** | **3** | **4** | **5** | **6** | **7** | **8** | **9** |
| --- | --- | --- | --- | --- | --- | --- | --- | --- | --- |
| Observed counts | 35 | 15 | 9 | 11 | 7 | 5 | 4 | 6 | 4 |
| NBL Expected Counts | 28.9 | 16.9 | 12.0 | 9.3 | 7.6 | 6.4 | 5.6 | 4.9 | 4.4 |
| Pearson Residuals | 1.13 | 0.46 | 0.86 | 0.56 | 0.22 | 0.56 | 0.66 | 0.49 | 0.19 |
| Observed Probability % | 36.5 | 15.6 | 9.4 | 11.5 | 7.3 | 5.2 | 4.2 | 6.3 | 4.2 |
| Expected Probability % | 30.1 | 17.6 | 12.5 | 9.7 | 7.9 | 6.7 | 5.8 | 5.1 | 4.6 |

**Declining Post-1990 Salamander Counts**
| **First Digits** | **1** | **2** | **3** | **4** | **5** | **6** | **7** | **8** | **9** |
| --- | --- | --- | --- | --- | --- | --- | --- | --- | --- |
| Observed counts | 65 | 32 | 21 | 10 | 21 | 4 | 11 | 5 | 4 |
| NBL Expected Counts | 52.1 | 30.5 | 21.6 | 16.8 | 13.7 | 11.6 | 10 | 8.8 | 7.9 |
| Pearson Residuals | 1.79 | 0.28 | 0.13 | 1.65 | **1.97** | **2.23** | 0.31 | 1.29 | 1.39 |
| Observed Probability % | 37.6 | 18.5 | 12.1 | 5.8 | 12.1 | 2.3 | 6.4 | 2.9 | 2.3 |
| Expected Probability % | 30.1 | 17.6 | 12.5 | 9.7 | 7.9 | 6.7 | 5.8 | 5.1 | 4.6 |

FSD sample size N = 173. Max Pearson Residual >1.96; Euclidean distance d\*n = 1.437 (p>0.05); Cohen-W = 0.328 (95% CI: 0.261-0.394); Kossovsky SSD = 119.4

**Appendix Figure 3.**
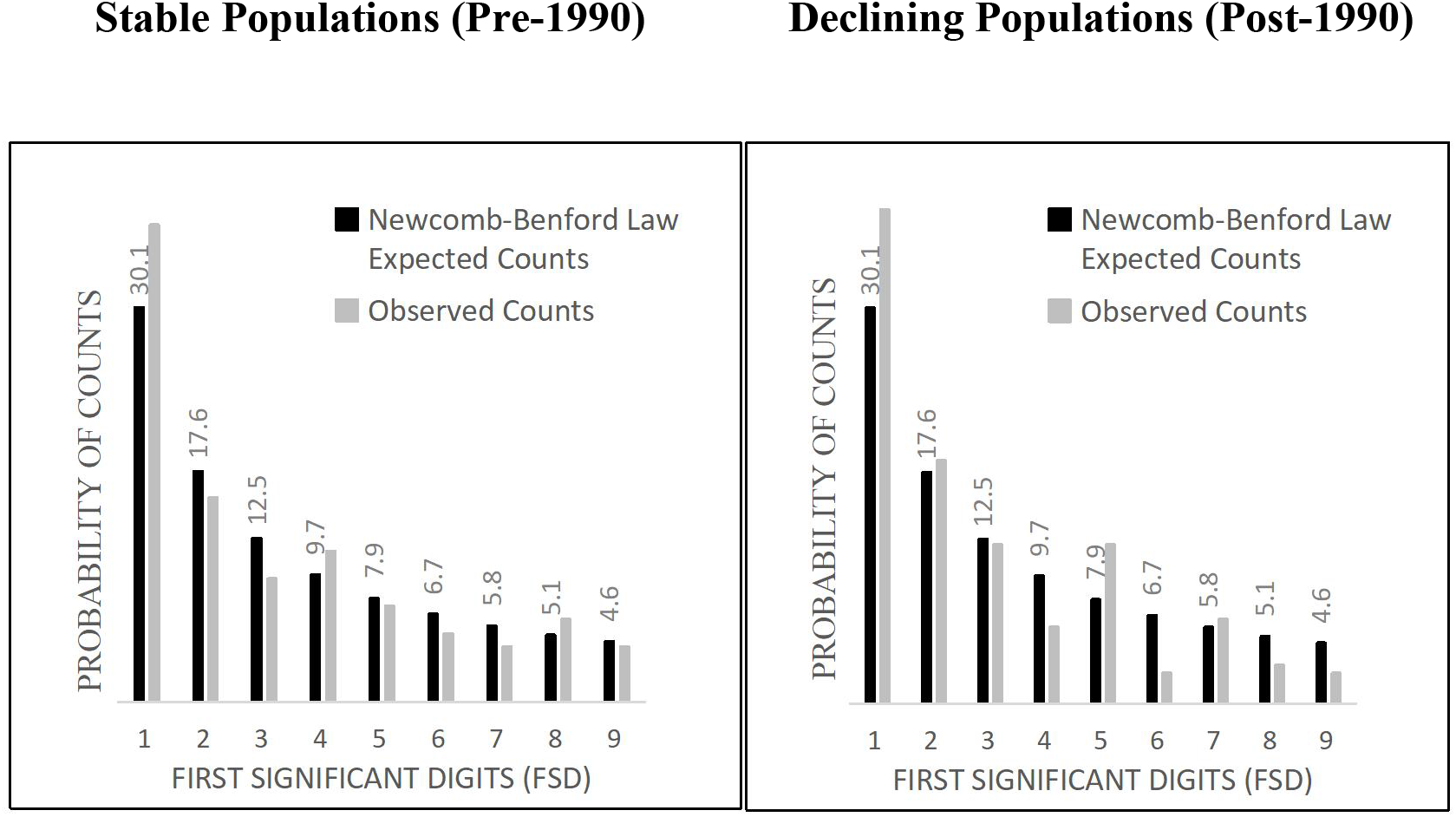
Comparison between FSD counts expected by Newcomb-Benford law and the probability distribution of observed counts for mean observations/site visit of multiple *Plethodon* salamander populations from Eastern United States [76]. Graphs of data from Appendix Table 3.

### Case Report 4: Regional fish communities with stable biotic integrity

#### Background

For the past six decades the State of Ohio Environmental Protection Agency (Ohio EPA) has conducted surveys of fish communities from streams and rivers to determine attainment of Federal Clean Water Act goals. Fish are sampled using standardized electrofishing protocols [79]. The overall condition and health of fish communities is evaluated using a modified Index of Biotic Integrity (IBI) [80], and agency developed Modified Index of Well-Being (MIwb). These indices are combined with information for benthic macroinvertebrate communities to assign Clean Water Act designated uses for protection of aquatic life. Stream segments designated Exceptional Warmwater Habitat (EWH) have unique species of fish with persistent levels of biotic integrity over time, and are associated with watersheds that have minimal background levels of anthropogenic disturbance. Chemical and biological survey results are available from Ohio EPA online technical documents:

[https://epa.ohio.gov/divisions-and-offices/surface-water/reports-data/biological-and-water-quality-reports].

Five EWH stream segments from various ecoregions of Ohio known to the author were selected to investigate correspondence of biomass measurements (reported as relative weight) with FSD predictions of the Benford distribution. Details of sample locations are as follows (sample date, river mile, # fish spp., # individual fish captured, length of stream sampled): Grand River (2004, 8.5, 26, 301, 0.42 km); Chagrin River (2003, 36.4, 21, 361, 0.2 km); Big Darby Creek (2014, 30.2, 39, 1012, 0.5 km); Scioto River (2011, 9.1, 33, 519, 0.5 km); Mohican River (2007, 6.5, 29, 360, 0.5 km).

Fish biomass measurements meet data collection constraints required to test NBL predictions. Fish were randomly sampled from all available habitats (riffles, run, pools) and the set of data spans three orders of magnitude, with total sample size sufficient for statistical inference when data from five EWH stream distributions are combined. Because raw data are scattered in multiple online technical support documents, biomass measures for 141 fish species from the five stream locations are given in Appendix Table 4, sorted by Benford FSD categories. This format of data presentation allows for calculation of biomass mean and variance for each of the nine FSD categories. As a point of reference, the 80.82 value in Appendix Table 4 represents the relative weight (kg/km) of 156 Golden Redhorse (*Moxostoma erythrurum*) collected from the Big Darby Creek, which represents ∼27.5% of total fish community biomass at that stream location.

#### Results and discussion

FSD biomass counts for 141 fish species collected from five EWH stream segments provide insufficient analytic weight of evidence to reject the tested (H_O_) of similarity between observed *vs* expected FSD probability distributions (Appendix Table 4 & Figure 4). Maximum Pearson Residuals (<1.96) and Cohen-W effect size magnitude (W=0.182) agree with results from statistical inference (d*n=0.759) that distributions of EWH fish communities over a wide geographic area of Ohio persist within a threshold range of stable dynamic equilibrium.

Odum [81] defined an ecological system as a set of interacting components forming a unified whole at the population, community, and ecosystem levels of organization. The geographically isolated distributions of EWH fish communities from this case report, combined with observations of widespread stable populations of salamanders in the eastern United States (case report #3), are consistent with the meta-community and meta-population concepts of organization proposed by Leibold et al. [82]. Benford FSD predictions of stability of meta-distributions are maximized at sample locations where constraint violations related to collection of data are minimized.

Taylor’s power law (V = aM^b^), where V = variance, M = mean, a = constant, and b=correlation slope, is an empirical observation relating the variance of measurements on organisms to mean unit area of sampling effort [62]. Fish species community biomass measurements from EWH stream segments reveal high coefficient of determination (Y = 3.1X ^2.1^, R^2^ = 0.91) between Taylor’s power law and the mean (X) and variance (Y) of data for the nine Benford FSD categories (Appendix Table 4). This correlation leads to a hypothesis that the FSD sequence of the Newcomb-Benford law provides an underlying mathematical explanation of the correlation slope metric of Taylor’s Law for different levels of ecological organization and spatial scale [62].

**Appendix Table 4.** Biomass (kg/km) of 141 species of fish collected from five streams in Ohio designated Exceptional Warmwater Habitat (EWH) for protection of aquatic life. Measurements sorted by Benford first digit categories.

| First Digits | 1 | 1 | 2 | 3 | 4 | 5 | 6 | 7 | 8 | 9 |
| --- | --- | --- | --- | --- | --- | --- | --- | --- | --- | --- |
|  | 0.01 | 0.19 | 0.02 | 0.03 | 0.04 | 0.05 | 0.06 | 0.07 | 0.08 | 0.09 |
|  | 0.01 | 1.06 | 0.02 | 0.03 | 0.04 | 0.05 | 0.06 | 0.73 | 0.08 | 0.09 |
|  | 0.01 | 1.08 | 0.02 | 0.03 | 0.04 | 0.05 | 0.6 | 0.76 | 0.87 | 0.9 |
|  | 0.01 | 1.1 | 0.02 | 0.03 | 0.04 | 0.56 | 0.63 | 70.18 | 80.82 | 0.99 |
|  | 0.01 | 1.2 | 0.02 | 0.03 | 0.04 | 0.56 | 0.64 |  |  | 9.0 |
|  | 0.01 | 1.2 | 0.02 | 0.03 | 0.04 | 5.10 | 0.64 |  |  | 9.31 |
|  | 0.01 | 1.52 | 0.02 | 0.03 | 0.40 | 5.20 | 0.69 |  |  |  |
|  | 0.01 | 1.68 | 0.02 | 0.32 | 0.46 | 5.51 | 6.30 |  |  |  |
|  | 0.01 | 1.71 | 0.02 | 0.34 | 0.48 | 5.67 | 6.7 |  |  |  |
|  | 0.01 | 1.98 | 0.02 | 3.2 | 4.0 | 53.85 | 62.25 |  |  |  |
|  | 0.01 | 10.52 | 0.2 | 3.2 | 4.01 |  | 67.5 |  |  |  |
|  | 0.01 | 11.03 | 0.2 | 3.6 | 4.8 |  |  |  |  |  |
|  | 0.10 | 12.0 | 0.21 | 3.68 | 44.90 |  |  |  |  |  |
|  | 0.11 | 14.11 | 0.29 | 3.7 | 48.17 |  |  |  |  |  |
|  | 0.11 | 14.72 | 2.0 | 32.99 |  |  |  |  |  |  |
|  | 0.11 | 15.8 | 2.0 | 33.6 |  |  |  |  |  |  |
|  | 0.12 | 16.94 | 2.07 | 34.3 |  |  |  |  |  |  |
|  | 0.14 | 18.22 | 2.23 | 38.03 |  |  |  |  |  |  |
|  | 0.14 | 19.5 | 2.25 |  |  |  |  |  |  |  |
|  | 0.14 | 114.5 | 2.41 |  |  |  |  |  |  |  |
|  | 0.15 |  | 2.49 |  |  |  |  |  |  |  |
|  | 0.16 |  | 2.8 |  |  |  |  |  |  |  |
|  | 0.16 |  | 2.80 |  |  |  |  |  |  |  |
|  | 0.16 |  | 2.85 |  |  |  |  |  |  |  |
|  | 0.17 |  | 21.01 |  |  |  |  |  |  |  |
|  | 0.17 |  | 21.30 |  |  |  |  |  |  |  |
|  | 0.18 |  | 21.35 |  |  |  |  |  |  |  |

| First Digits | 1 | 2 | 3 | 4 | 5 | 6 | 7 | 8 | 9 |
| --- | --- | --- | --- | --- | --- | --- | --- | --- | --- |
| Observed Counts | 47.0 | 27.0 | 18.0 | 14.0 | 10.0 | 11.0 | 4.0 | 4.0 | 6.0 |
| NBL Expected Counts | 42.4 | 24.8 | 17.6 | 13.7 | 11.1 | 9.4 | 8.2 | 7.2 | 6.5 |
| Pearson Residuals | 0.71 | 0.44 | 0.09 | 0.08 | 0.33 | 0.52 | 1.47 | 1.19 | 0.20 |
| Observed % Probability | 33.3 | 19.1 | 12.8 | 9.9 | 7.1 | 7.8 | 2.8 | 2.8 | 4.3 |
| Expected % Probability | 30.1 | 17.6 | 12.5 | 9.7 | 7.9 | 6.7 | 5.8 | 5.1 | 4.6 |
| Digit Mean (X) | 5.94 | 3.28 | 8.74 | 7.68 | 7.67 | 13.27 | 17.93 | 20.46 | 3.40 |
| Digit Variance (Y) | 301 | 43 | 208 | 274 | 270 | 658 | 1213 | 1619 | 20 |

FSD sample size N = 141. Max Pearson Residual<1.96; Euclidean distance $d^*n = 0.638$ ( $p>0.10$ ); Cohen-W = 0.182 (95% CI: 0.134- 0.230); Kossovsky SSD = 28.9. Taylor's Power Law: $Y = 3.1 X^{2.1}$ , $R^2 = 0.91$ .

**Appendix Figure 4.**
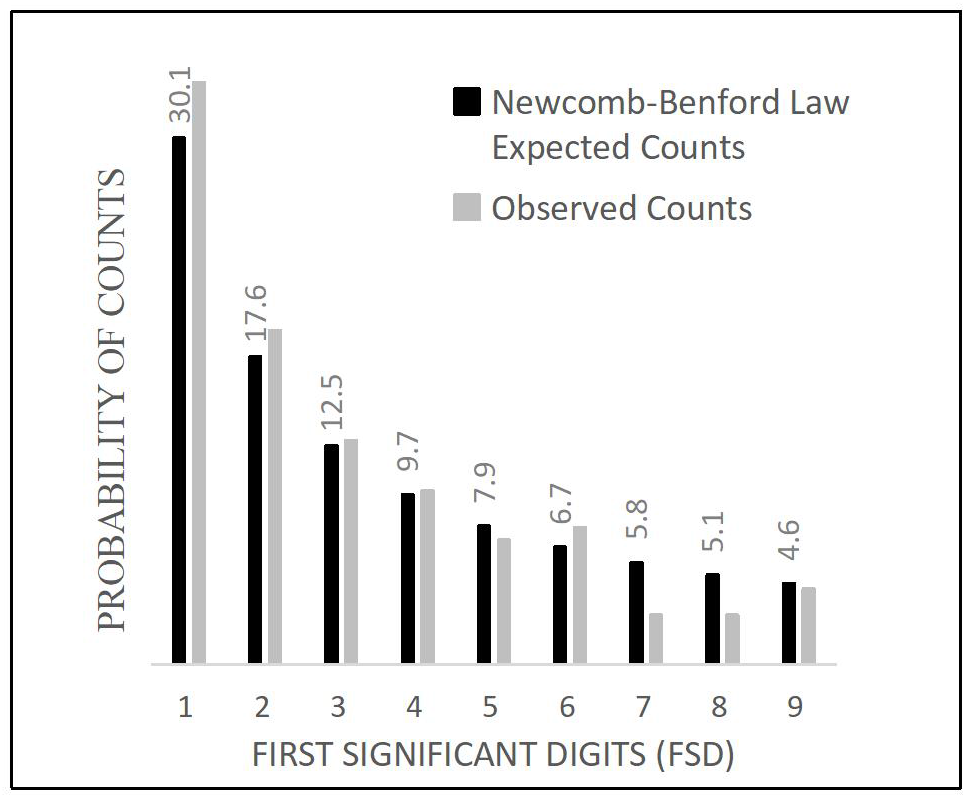
Comparison between FSD biomass counts expected by Newcomb-Benford law and the probability distribution of observed counts for 141 species of fish collected from stream segments with persistent exceptional biotic integrity. Graph of data from Appendix Table 4.

